# Characterizing mixed single chain amphiphile-based coacervates as a robust protocell system

**DOI:** 10.1101/2025.04.05.647343

**Authors:** Gauri M. Patki, Vanthanaa Sridhar, Sudha Rajamani

## Abstract

Prebiotic soup would have been a dilute pool of various constituent chemicals that would have reacted with each other to form biologically relevant precursors during life’s origin. In this milieu, compartments formed by liquid-liquid phase separation (LLPS) are thought to have facilitated concentration of chemicals, thereby catalyzing their reactions. Towards this, various LLPS-based systems have been studied as model protocells. Relevantly, fatty acid-based (decanoic acid) coacervates have recently been explored as model protocells. As far as protocell research is concerned, fatty acids have been studied much more extensively in the context of vesicle-forming entities when compared to them resulting in coacervate systems. Furthermore, exogenous delivery and endogenous synthesis of fatty acids suggest the prevalence of single chain amphiphiles (SCAs) on the early Earth, with a greater abundance of the shorter chain length moieties. In this backdrop, we set out to fabricate robust coacervate-based protocells using SCAs that would have been readily present in a chemically heterogeneous prebiotic soup, and which could thrive under various prebiotically relevant selection pressures. Towards this, we characterized a mixed amphiphile-based coacervate system composed of nonanoic acid (NA), nonanol (NOH) and tyramine (Tyra), which could form coacervates over a broad range of pHs, temperatures, and salt concentrations. This is noteworthy as compositionally heterogenous vesicles have also been shown to have advantages over pure fatty acid vesicles. Additionally, we also demonstrate RNA sequestration in these coacervates that gets enhanced upon addition of cationic amino acids, emphasizing the importance of co-solute interactions in the prebiotic soup. Lastly, we also demonstrate nonenzymatic template-directed primer extension in these coacervates, suggesting the potential functional role of these compartments during life’s origin.

## 1. Introduction

Modern living cells are comprised of a humongous framework of biomolecules and biochemical reactions that are spatially concentrated in different regions by a fundamental principle of compartmentalization.^[1,2]^ Although spatially segregated, these compartments perform biochemical reactions in a harmonious and consistent manner so as to produce energy and raw materials that are required for the cell to undergo growth and division. These complex entities are considered a result of evolution from much simpler compartmentalized systems, which are considered to have existed on the prebiotic Earth.^[3,4]^ These primordial compartments known as ‘protocells’ are hypothesized to have formed in the prebiotic soup. They would have possessed the ability to concentrate monomeric molecules and oligomers like nucleic acids/proteins, while also facilitating prebiotically plausible reactions and undergoing growth and division.^[5–7]^ In this regard, study of prebiotically plausible protocells has mainly involved the characterization of two typed of model systems – membrane bound fatty acid-based vesicles and membraneless compartments that are formed by liquid-liquid phase separation (LLPS).

The reasons why fatty acid vesicles have been extensively studied as model protocells are multifold^[8–13]^. First, small chain fatty acids (SCFAs) are thought to have been exogenously delivered on the early Earth^[14– 17]^ and their terrestrial synthesis under prebiotically plausible conditions has also been demonstrated.^[18,19]^ Second, they form vesicles spontaneously when the pH of the surrounding environment is equivalent to the pKa of their carboxylic acid head group.^[12,14]^ Third, they have the potential to encapsulate a range of solutes, including small molecules, nucleic acids, proteins, and the capacity to facilitate related prebiotically pertinent reactions such as nonenzymatic oligomerization/replication and ribozyme catalysis, etc.^[7,17,20–23]^ These are the properties that also make them resemble today’s cellular compartments. However, pure fatty acid vesicles are stable only over a narrow pH range. In order to increase the stability of these vesicles over a wide pH range, mixed amphiphile systems have been characterized extensively.^[13,24,25]^ Specifically, doping the pure fatty acid system with alcohol and glycerol monoester containing derivatives has been shown to substantially increase the stability of the vesicles across a broad range of pH and salt concentrations.^[13,24]^ Despite these characteristics of fatty acid-based vesicles that make it close-to-ideal as a model protocell, the bilayer membrane acts as a semipermeable barrier to the diffusion of large and polar molecules across the membrane. Conversely, compartments formed by LLPS, including aqueous two-phase systems (ATPS) or coacervates, act as open systems that allow ready diffusion of solute molecules within the compartment as well as with the bulk solution.^[26–29]^ Further, LLPS compartments have been shown to sequester biomolecules, and consequently increase their effective concentration within these compartments by several orders of magnitude.^[30–32]^. Additionally, they have also been demonstrated to increase the rate of ribozyme-based as well as metabolic reactions.^[29,31–36]^

With an aim to explore such ‘open’ systems as protocell models, several LLPS systems have been developed and studied over the last decade.^[29]^ Recently, a myristic acid-guanidinium hydrochloride-based system was described, where the myristic acid was shown to reversibly transition from vesicles (at pH 8.5) to coacervates (at pH 9.5). This was shown to result from the interaction of guanidinium ions with higher order aggregates of micelles, which form at alkaline pH.^[37,38]^ Even though this system seems prebiotically relevant; the resultant coacervates were shown to sequester RNA poorly. Furthermore, RNA is prone to getting hydrolyzed readily at pH of 9.5 where the coacervates actually form. This problem of alkaline pH was partially circumvented by the demonstration of a coacervate system that used decanoic acid vesicles, which were shown to transition into coacervate droplets upon addition of dopamine.^[39]^ However, dopamine is known to form polydopamine as a result of spontaneous oxidation, which led to the collapse of the coacervate upon increasing the pH beyond 7.5.^[39]^ Furthermore, it was shown to sequester hydrophobic and cationic molecules by 1-2 orders of magnitude when compared to anionic molecules, thus rendering the system less efficient for the sequestration of RNA.^[39]^ Moreover, the system was found to be stable for only a short time (∼1 hr) after which it started forming crystals.^[40]^

In the aforementioned backdrop, we set out to characterize short chain fatty acids that could not only form coacervates but also sequester RNA; an important informational molecule of the primordial RNA World. In addition to being prebiotically relevant when compared to systems comprising of polymer-based coacervates (such as PDDA-spermine), short chain fatty acids are hypothesized to be prebiotically more abundant than the longer chain fatty acids.^[16,18,19]^ Nonetheless, they are much less explored than longer chain fatty acids as compartmentalization systems because those that can form vesicles (especially C8-10 chain length SCAs), result in very dynamic and, consequently, leaky membranes. In this study, we characterized a C9-based mixed amphiphile coacervate system. We demonstrate how these nonanoic acid (NA) and nonanol (NOH) containing vesicles transition into coacervates upon the addition of tyramine (Tyra). The resultant NA-NOH/Tyra coacervates undergo coalescence, a quintessential property of LLPS; while also readily forming droplets upon mixing even after several days, unlike the previously described system (decanoic acid-dopamine). Furthermore, they were found to be stable across pHs (from pH 7 to 9), and temperatures ranging from 5°C to 65°C. Pertinently, they also could sequester biomolecules such as protein and RNA. To the best of our knowledge, we report for the first time a mixed amphiphile-based coacervate system, which can tolerate various prebiotically pertinent selection pressures as well as sequester biomolecules like RNA, making it a robust protocellular model than the pure fatty acid-based coacervates. In all, these studies underline how the compositional heterogeneity of the prebiotic soup could have enabled the formation of robust SCA-based coacervates; entities that were minimally discussed in the context of vesicle systems. Further, such studies enable describing amphiphilic moieties that can eventually be used in fabricating multi-compartment cells, setting the stage for characterizing more extant cell-like entities wherein physicochemically distinct LLPS compartments could be used to facilitate different kinds of reactions within the same protocell.

## 2. Results

### 2.1. A mixed amphiphile-based coacervate system as a prebiotic model protocell

With an intention to discern short chain fatty acids other than decanoic acid that can form coacervates, we first checked whether dopamine (DPA) could induce vesicle to coacervate transition in pure nonanoic acid (NA) vesicles. We used dopamine, because it was earlier reported to induce vesicle to coacervate transition in decanoic acid (DA) vesicles. When dopamine was added to nonanoic acid vesicles at pH 7, an increase in turbidity as observed in the sample, which upon visualizing under microscope, showed droplets, as reported for DA vesicles.^[39]^ The stability of the droplets was evaluated across different pHs (7, 8, and 9) since prebiotic pools are thought to have had variable pH because of their diverse constituents.^[25,41]^ Droplets were observed only at pH 7 and this can be attributed to the absence of nonanoic acid vesicles at pHs above 7 (Figure 1a, upper panel). To increase the pH range across which coacervate droplets could be observed, pure nonanoic acid was doped with nonanol (NOH) in 10:1 ratio to prepare NA-NOH mixed vesicles (S1). Coacervation requires vesicles to exist in the first place and NA-NOH system has been shown to form vesicles across a broad range of pHs.^[24]^ Relevantly, with the NA-NOH/DPA system, we observed coacervate droplets at all the three pHs (Figure 1a, lower panel). A protocellular system that can tolerate a broad range of pHs would have had higher chances of survival in the environment of changing pH on the early Earth. Thus, the mixed amphiphile-based coacervate system would have been beneficial over the pure system under related selection pressures. In this regard, we wish to highlight that although the importance of compositional heterogeneity in the formation of robust protocells has been established for fatty acid vesicles, it is largely unexplored in the context of coacervate-based systems.^[6,13,25,42,43]^

**Figure 1:**
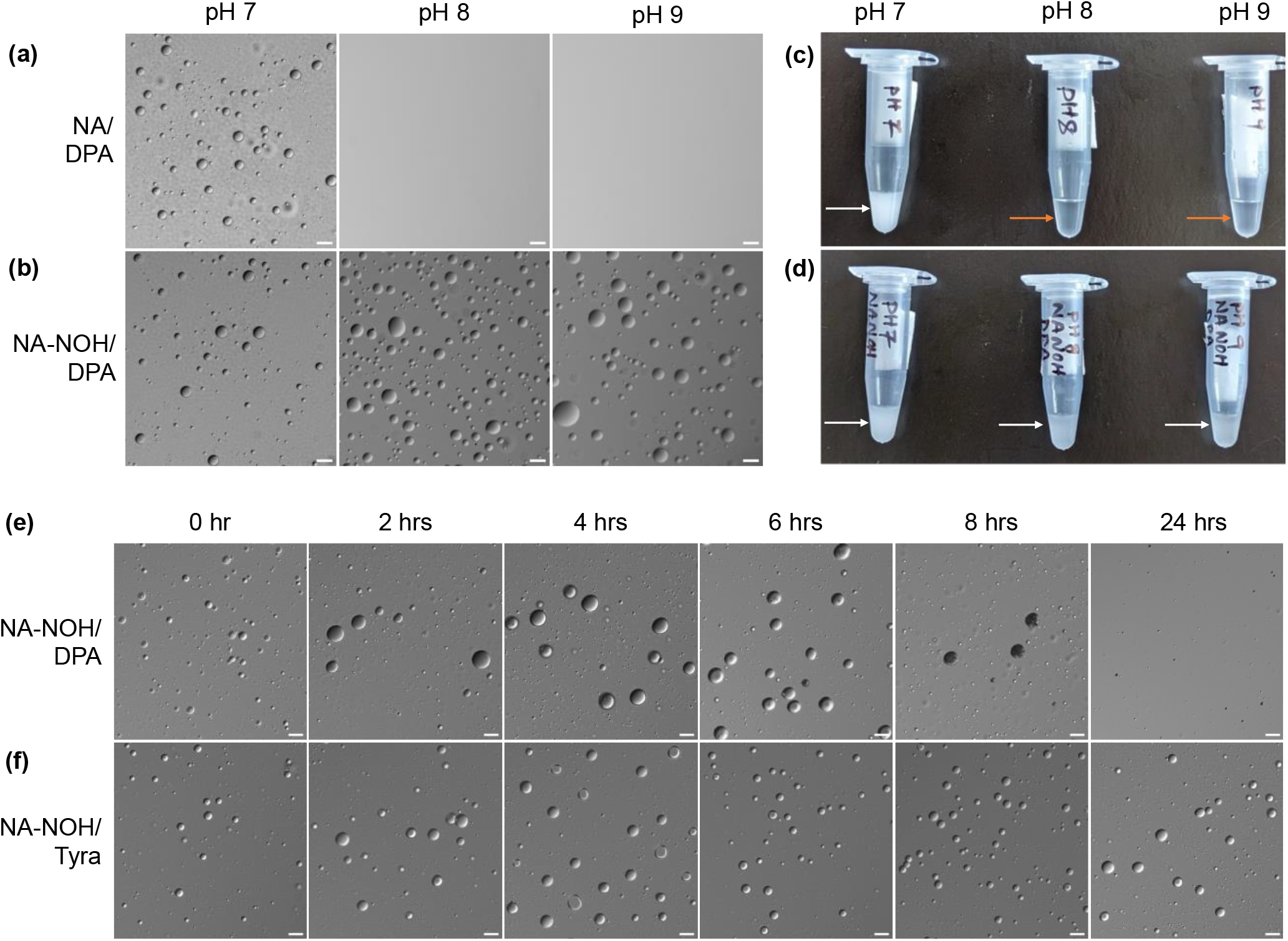
Compositional heterogeneity forms robust coacervate droplets. Pure NA/DPA based coacervates forming only at pH 7 **(a)**, and the reflection of their presence on the turbidity of the samples **(c). (b)** Compositionally diverse, mixed NA-NOH/DPA based coacervates forming across a broad pH range (7 to 9), and the reflection of their presence on the turbidity of the samples **(d)**. The white and the orange arrows in **(c)** and **(d)** represent turbid samples and clear samples, respectively. It was observed that the NA-NOH/DPA sample started to turn darker with time, and the droplets could not be observed after 6 hours **(e)**. To increase the stability and the robustness of the system, DPA was replaced with Tyra as it does not undergo spontaneous oxidation like DPA does. Indeed, the NA-NOH/Tyra system continued to form coacervate droplets over a much longer time (24 hours) **(f)** than NA-NOH/DPA system. Scale bar: 20 μm. N=3.

### 2.2. Towards constructing robust NA-NOH-based coacervates: Replacing dopamine with tyramine

Dopamine is known to undergo spontaneous oxidation to form polydopamine (melanin). Hence, it could be hypothesized that when dopamine is used for inducing coacervation, it would get consumed over the course of time in the absence of any antioxidizing agent. In order to check whether the NA-NOH/DPA droplets could persist even after the spontaneous oxidation of dopamine, we prepared NA-NOH/DPA droplets in the absence of any antioxidizing agent and observed the presence of droplets over time. We observed that the sample progressively turned brown with time (Figure S2). Additionally, on microscopic analysis, it was qualitatively observed that the number of droplets went down with time and no droplets could be observed after 8 hours (Figure 1e). This indicated that the consumption of dopamine through its spontaneous oxidation was resulting in the reduction in the number of NA-NOH/DPA droplets with time possibly due to their disruption.

To overcome the issue of spontaneous oxidation and form a more robust and durable coacervate system, dopamine was replaced with tyramine (Tyra) to induce vesicle-to-coacervate transition in NA-NOH vesicles. In stark contrast to NA-NOH/DPA droplets, NA-NOH/Tyra droplets could be observed even after days (Figure 1f, S3). In both the systems, the droplets would coalesce in the tube with time; however, they would reappear upon gentle mixing, making the suspension turbid. This reappearance of droplets after mixing was seen only for the first six hours in NA-NOH/DPA system, whereas, for NA-NOH/Tyra system, the same was observed even after a few days (S3). Further, at no point did the NA-NOH/Tyra system turn brown, highlighting the robustness of the NA-NOH/Tyra system over the one composed of NA-NOH/DPA. Having established that this new NA-NOH/Tyra system was indeed a more robust coacervate system, we next took a systematic approach to characterize the system for its potential to be acknowledged and used as a model protocell.

Towards this, it was essential to first determine the range of concentrations of NA-NOH and tyramine that resulted in the formation of droplets. It has been reported earlier that the critical vesicle concentration (CVC) of NA-NOH (10:1) system is 20 mM. Since it was established earlier that the presence of NA-NOH vesicles is indeed required for coacervates to form (Figure 1a-d), we chose to stay well above this CVC and worked with total amphiphile concentrations of 40, 50, and 60 mM (Figure S4), where the NA-NOH vesicles would surely be present. Next, for every NA-NOH concentration, a range of Tyra concentrations were scanned, beginning with a concentration that was equivalent to that of NA-NOH in the system. A general trend was observed in which the size of the droplets increased upon increasing the concentration of NA-NOH. The spike in the turbidity indicates the transition of the vesicles into droplets (Figure S5). In this context, the higher error bars seen in case of 50/80 and 60/70 NA-NOH/Tyra systems (Figure S4) may have been because of the different optical properties of these suspensions since they contained a co-existing population of droplets and other higher order aggregates (Figure S5). And, at these ratios, there could be transition occurring from vesicles to coacervates.

Based on microscopic observation, we narrowed down to NA-NOH concentration of 60 mM for all further characterization because it yielded consistent and numerous droplets at and above 80 mM Tyra concentration (Figure S5). To further understand the coacervation limit of this system, Tyra concentrations ranging from 60 mM to 350 mM were scanned (Figure 2a). It was found that at all the Tyra concentrations that were studied (from 80 mM to 200 mM), droplets were observed (Figures S5-S6). Furthermore, from visual analysis of microscopy data, it was seen that the size of the droplets was found to increase with increasing Tyra concentration, especially from 140 mM onwards. At 250mM, 300 mM and 350mM Tyra, very few but huge droplets were observed (Figure S6). These results suggested that for 60 mM NA-NOH vesicles, coacervates could be observed for a very broad range of Tyra concentrations. Furthermore, we evaluated the micropolarity of the coacervates as a proxy to understand whether the coacervates formed using the different tyramine concentrations, resulted in coacervates with similar internal environment. Towards this, Laurdan was used as a solvatochromic probe.^[44–46]^ and as a first step, the partitioning of Laurdan into the coacervates was checked (Figure S7). After ensuring that the emission was being recorded only in the coacervate phase, we went ahead and calculated the generalized polarization (GP) values. The GP value of NA-NOH vesicles increased upon adding tyramine to the vesicles, suggesting that the micropolarity of the coacervates is significantly higher than that of the vesicles (Figure 2b). Furthermore, statistically non-significant difference was observed across coacervates containing various tyramine concentrations ranging from 80 mM to 350 mM, suggesting that they could have similar microenvironments. The observation that coacervates could be formed at various Tyra concentrations is noteworthy, as there is a plausibility that Tyra or Tyra-like molecules, which can induce vesicle to coacervate transition, may have been available at variable concentrations in the prebiotic soup.

**Figure 2:**
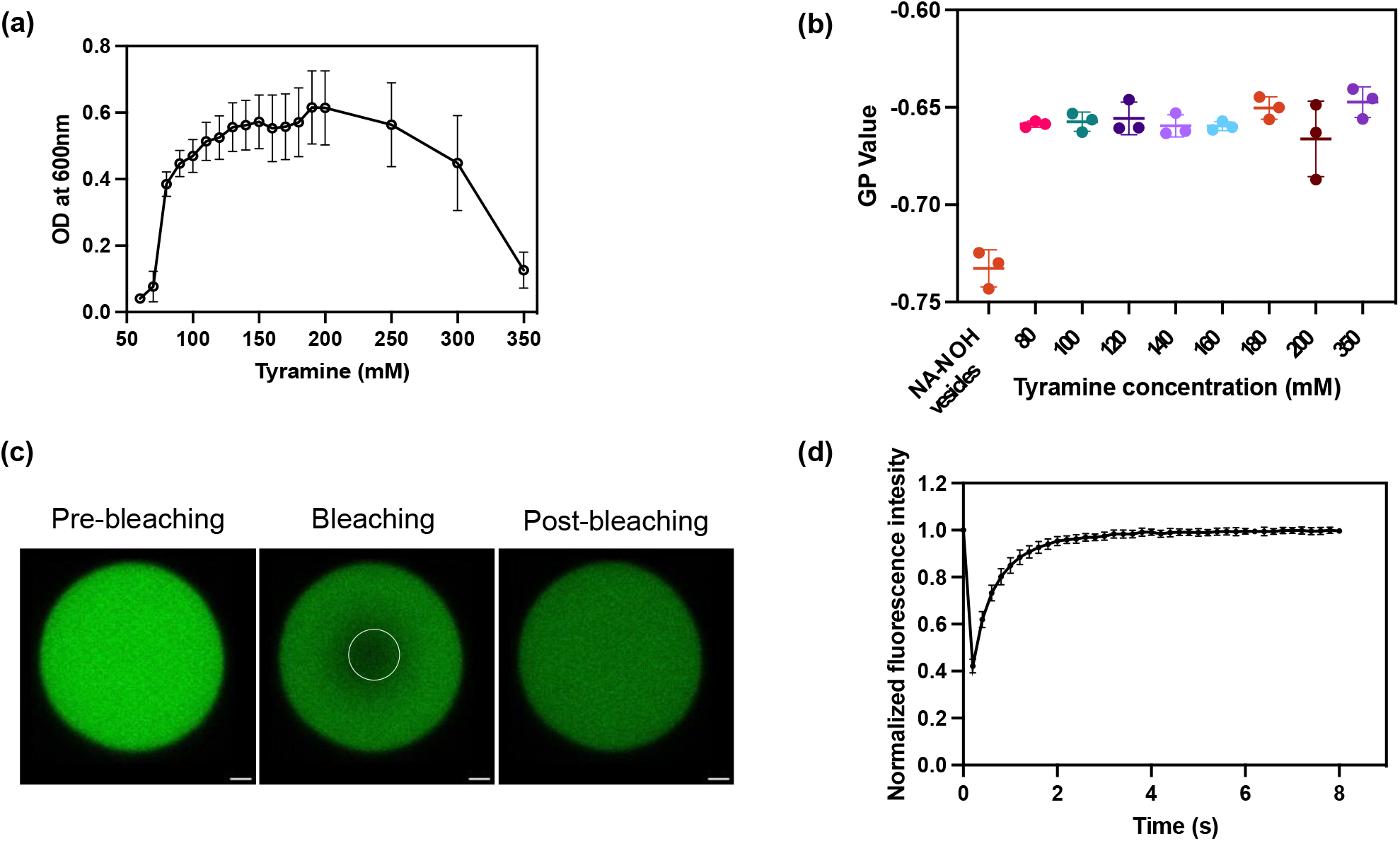
Characterizing NA-NOH/Tyra system as model protocell. **(a)** Scanning the concentrations of tyramine across which 60 mM NA-NOH forms droplets. **(b)** Laurdan GP values of coacervates prepared from 60 mM NA-NOH with variable tyramine concentrations. We performed Tuckey’s multiple comparisons test on this dataset, where we observed statistically nonsignificant difference between the coacervate samples containing tyramine concentrations ranging from 80 mM to 350 mM (p > 0.5). All the populations containing tyramine, however, were found to be statistically significantly different from the corresponding vesicle population (p < 0.0001). **(c)** To understand whether the NA-NOH/Tyra droplets allow molecular diffusion, 0.5 μM GFP was sequestered within the droplets. Panel **(c)** represents sequestered GFP (pre-bleaching) and partial photobleaching (white circle) of the sequestered GFP followed by fluorescence recovery within the bleached spot (post-bleaching). **(d)** Quantification of FRAP experiments. Error bars: Standard deviation. N=3; n=30. Scale bar: 1μm.

### 2.3. NA-NOH/Tyra droplets enable molecular diffusion within the compartments

Life is thought to have emerged in a chemically heterogenous prebiotic soup, where interactions between the reactants would have resulted in the formation of precursors of biomolecules, including nucleosides, nucleotides, amino acids etc. These, in turn, would have interacted with each other to form more complex biomolecules like nucleic acids, peptides etc.^[47]^ However, since the prebiotic soup is thought to have been dilute, mechanisms to concentrate these molecules would have been crucial to facilitate their interactions. In this regard, LLPS compartments have been demonstrated to sequester biomolecules and increase their local concentration by several orders of magnitude than that in the bulk solution, as well as to enhance the rates of the reactions being contained within them.^[29,30,32]^ For molecular reactions to take place inside these compartments, the coacervates should facilitate diffusion of molecules within the interior. Given this, we checked whether the NA-NOH/Tyra droplets can allow molecular diffusion. For this, GFP was sequestered within the droplets and subjected to partial photobleaching, and the fluorescence recovery after photobleaching (FRAP) was monitored.

As is evident in Figures 2c and 2d, the photobleached spot recovered fluorescence, confirming that the droplets allow for molecular diffusion, while also corroborating that the NA-NOH/Tyra droplets are indeed LLPS compartments. The half-time of recovery of fluorescence and the apparent diffusion coefficient of GFP in 60/100 NA-NOH/Tyra system were found to be 0.41 ± 0.07 s and 0.14 ± 0.02 μm^2^s^-1^, respectively. Furthermore, this implied that such systems could facilitate prebiotically plausible reactions between the molecules sequestered within the compartments since it allows molecular diffusion. Additionally, it would also support the exchange of molecules with the bulk solution; a process central for taking up fresh nutrients inside these protocells while also allowing for the removal of waste material out of the system. Thus, fatty acid coacervates provide an advantage over fatty acid vesicles as model protocells, wherein similar kind of ready exchange of molecules is limited by the bilayer of the fatty acid vesicles. In this backdrop, we characterized the tolerance of these systems to various selection pressures to further discern the robustness of NA-NOH/Tyra coacervates as model protocells.

### 2.4. NA-NOH/Tyra system tolerates prebiotically plausible selection pressures

Emergence of early life would have been shaped by ocean tides, hence protocellular assemblies would have required to survive selection pressures such as high salt concentrations, fluctuating pH and temperatures.^[48]^ Given this, the tolerance of 60/100 NA-NOH/Tyra system towards various concentrations of monovalent (NaCl) and divalent (MgCl_2_) cations was systematically estimated. As per the RNA World hypothesis, RNAs are thought to have played the dual role of acting as a genetic polymer while also catalyzing chemical reactions (ribozymes).^[49]^ For the ribozyme to catalyze chemical reactions, the presence of divalent cations, especially Mg^2+^, is extremely crucial. Such dual-function genetic polymers are thought to have been encapsulated inside protocells to result in functional cellular moieties[6]. However, fatty acid vesicles, which have been been demonstrated to encapsulate genetic polymers, are known to form precipitates of fatty acid salts at concentrations of divalent cations above 2-4 mM.^[20,50]^ Therefore, we first checked the stability of NA-NOH/Tyra system towards various MgCl_2_ concentrations. Figure 3a shows that the NA-NOH/Tyra droplets could be observed till 20 mM MgCl_2_ concentration, after which crystals were forming. Similarly, the stability of the droplets was checked over a range of NaCl concentrations since early ocean NaCl concentrations are thought to be as high as 600 mM.^[48]^ Figure 3b shows that NA-NOH/Tyra system forms droplets at a wide range of NaCl concentrations, ranging from 10 mM and all the way through 600 mM. However, visual observation suggested that the number of droplets reduced with increasing concentrations of NaCl, which was also corroborated by turbidity measurements (Figure S8).

**Figure 3:**
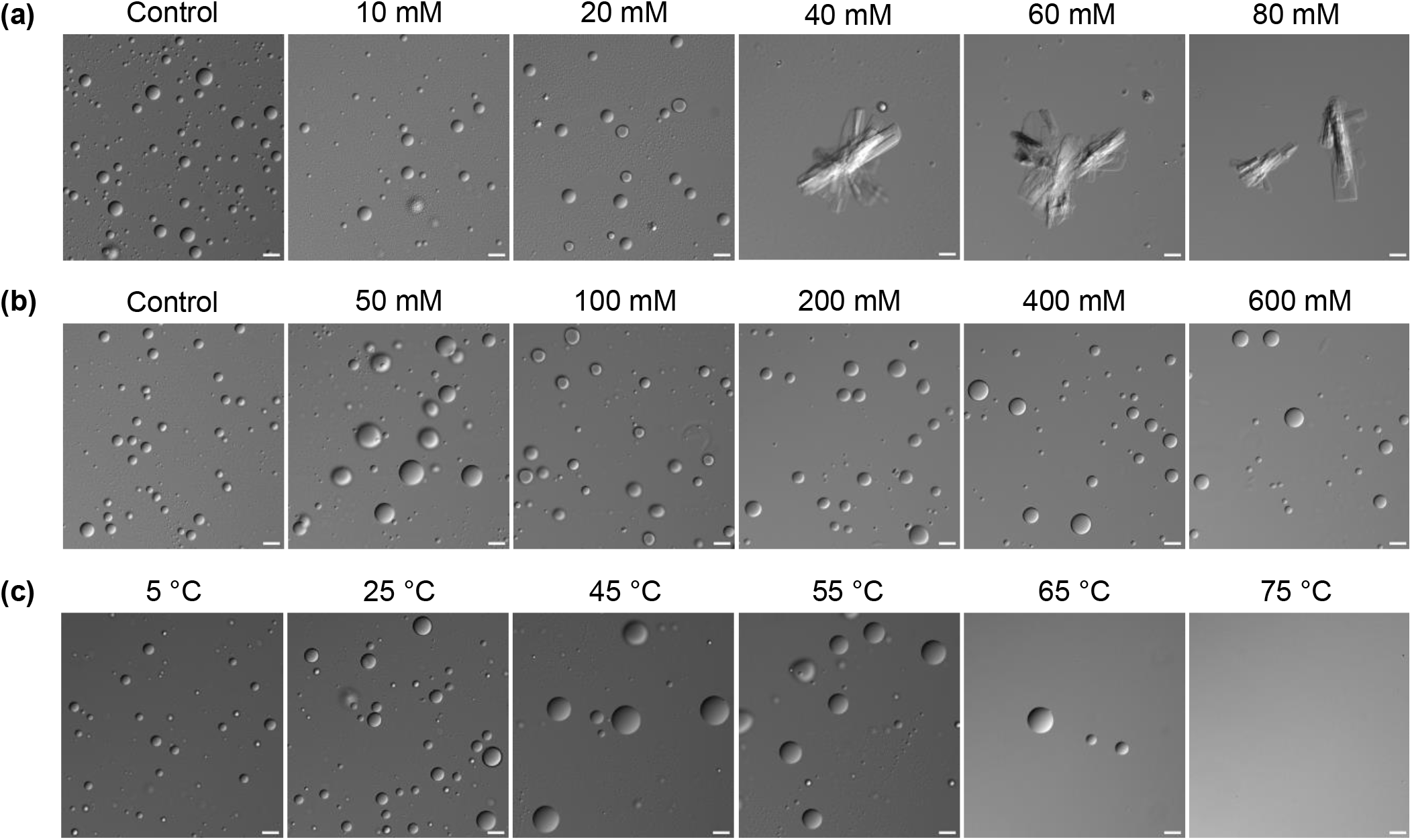
Tolerance of NA-NOH/Tyra system towards prebiotically pertinent selection pressures. Salt tolerance of 60/100 NA-NOH/Tyra droplets to various concentrations of MgCl_2_ **(a)** and NaCl **(b)**. When 60/100 NA-NOH/Tyra samples were incubated at various temperatures for 30 mins, ranging from 5°C to 75°C, droplets were observed till 65°C **(c)**. The density of the droplets, however, reduced substantially at 65°C. The sample incubated at 75°C for 30 mins appeared clear, and no droplets were observed. N=3. Scale bar: 20 μm.

After determining the tolerance of the NA-NOH/Tyra droplets to monovalent and divalent cations, we further subjected the droplets to various temperatures. Checking the stability of the droplets at various temperatures is important since temperatures on the prebiotic Earth are thought to have been variable.^[48,51,52]^ Notably, droplets were observed at all the temperatures tested, ranging from 5°C to 65°C; above this no droplets were present and the solution was clear (Figures 3c and S9). These results suggested that the NA-NOH/Tyra system has an upper critical solution temperature (UCST) of 65°C, which is the temperature above which the system does not phase separate to result in droplets. Overall, the observations in Figure 3 suggest that the NA-NOH/Tyra system is a robust one and shows tolerance to prebiotically relevant temperatures and high salt concentrations. In the context of MgCl_2_, although the coacervate droplets crashed out at concentrations above 20 mM, they could have still supported chemical reactions crucial to the putative RNA World hypothesis. These include nonenzymatic RNA replication and certain ribozyme-based reactions, which do not necessarily need concentrations of MgCl_2_ higher than 20 mM. As a first step towards undertaking such reactions within fatty acid coacervates, we asked whether RNA could get sequestered within the NA-NOH/Tyra droplets.

### 2.5. RNA sequestration inside the NA-NOH/Tyra droplets: Implications for nonenzymatic RNA replication

RNA has been shown to get sequestered within complex coacervates.^[29,32]^ Complex coacervates are a type of associative LLPS, wherein the interactions between two oppositely charged polyelectrolyte species results in their phase separation from the bulk solvent.^[29]^ In these cases, one of the two components forming the coacervates is a cationic polyelectrolyte, which can offer interactions with the anionic RNA molecules, and thus, can readily facilitate the sequestration of RNA inside the droplets.^[29,34,53,54]^ On the other hand, the two fatty acid coacervates studied to date have been reported to be inefficient in sequestering nucleic acid, mainly because of the repulsion between two anionic species, i.e. the fatty acids and the oligonucleotide. In order to circumvent this problem, cationic fatty acids like cetylpyridinium chloride and cetyltrimethylammonium bromide have been used to make coacervates, which were found to sequester DNA.^[55]^ However, the prebiotic relevance of these amphiphiles is unknown and possibly improbable.^[55]^

In this backdrop, we wanted to check whether our NA-NOH/Tyra droplets could sequester RNA. Towards this, we incorporated a Cy3 labelled 22-mer RNA and looked for Cy3 fluorescence inside the droplets. Figure 4a shows that the 60/100 NA-NOH/Tyra droplets do indeed show fluorescence, suggesting the sequestration of RNA within the droplets. However, while imaging the droplets, background fluorescence was also being observed suggesting that not all RNA was getting sequestered into the droplets. As reported previously, this could be because of the internal hydrophobic environment and is getting compounded due to repulsion coming from the negatively charged fatty acids constituting the droplets.^[39]^ Therefore, we wondered whether any simple, cationic molecules, which could have been present as co-solutes along with the droplets in the prebiotic soup, could enhance the sequestration of RNA. Towards this, cationic amino acids and prebiotically relevant co-solutes like lysine and arginine were evaluated for their role in RNA sequestration. Figure 4b shows that with the addition of lysine, RNA sequestration was significantly enhanced (Figure 4c). A similar effect was observed when arginine was added as a co-solute (Figure S10). Additionally, by doing Z stack analysis of the images, we also confirmed the even distribution of RNA across the droplet (Figure S11).

**Figure 4:**
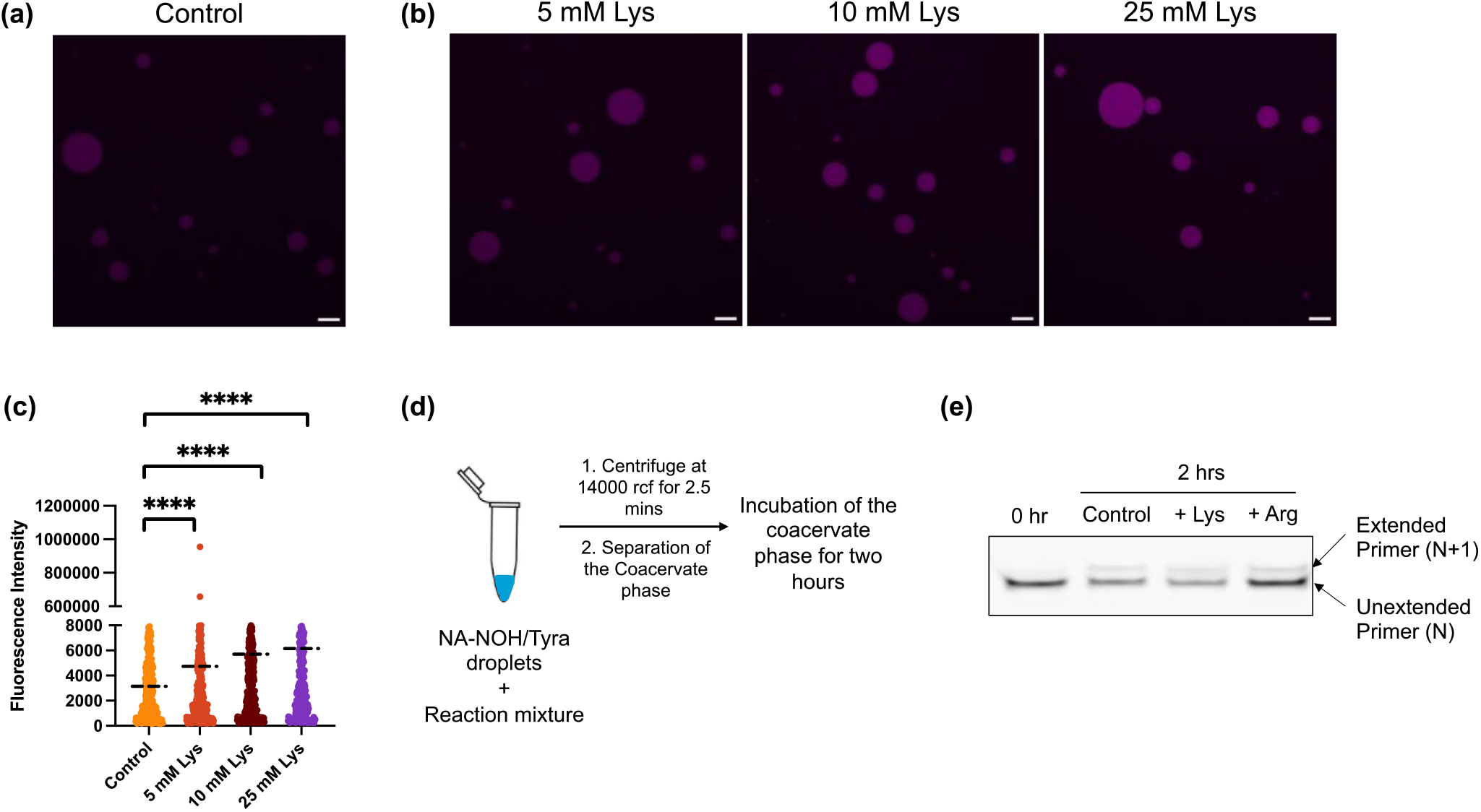
Effect of co-solutes in enhancing RNA sequestration. Confocal microscopy images showing the sequestration of Cy3 labelled RNA within 60/100 coacervate droplets **(a)** enhanced upon addition of cationic amino acid, Lysine, as a co-solute **(b)**. Scale bar: 10 μm. Quantification of the Cy3 fluorescence inside the droplets with varying concentrations of lysine **(c)**. Here, Kruskal-Wallis test was performed to statistically compare the fluorescence intensities between the droplets samples without the co-solute (control) and those with different concentrations of the co-solute. p<0.0001. N=3; n: at least 300 droplets per sample. Reaction scheme nonenzymatic template-directed replication of RNA in the 60/100 NA-NOH/Tyra coacervate phase **(d)**. Extension of the primer being observed when template-directed nonenzymatic replication reaction was allowed to proceed in the coacervate phase without amino acid (control), and with either lysine or arginine **(e)**.

Once RNA sequestration by NA-NOH/Tyra droplets was confirmed, we further asked whether they could facilitate nonenzymatic replication reactions. Towards this, an RNA primer-template complex was encapsulated inside the 60/100 NA-NOH/Tyra droplets, and a nonenzymatic replication reaction was initiated by adding imidazole activated nucleotides. Specifically, we studied the incorporation of adenosine monophosphate (AMP) on to the 3’ end of the template-bound primer against the templating base U, by using imidazole activated AMP (ImpA). To the encapsulated RNA primer-template complex, we added ImpA and immediately centrifuged the samples to separate the coacervate phase from the bulk solvent (Figure 4d). The separated coacervate phases from the individual reactions were incubated for 2 hours, which was followed by running them on 22% denaturing urea PAGE (Methods). Upon visualizing the gel, the extension of the primer was observed in all the reactions (Figure 4e). These reactions were performed in the presence of 25 mM Lysine and Arginine while control reactions without amino acids were also simultaneously carried out. To the best of our knowledge, this is the first ever demonstration of nonenzymatic template-directed RNA replication in a fatty acid-based coacervate system. In all, these results highlight the importance of chemical heterogeneity in the prebiotic soup and what it means for making robust and functional protocells of different types (i.e. membraneless and membrane-bound).

## 3. Discussion

Short chain fatty acids are thought to be prebiotically more abundant than long chain fatty acids.^[16,18,19]^ However, since the dynamicity of vesicles that result from short chain fatty acids is very high, their role in forming prebiotic compartments has remained less explored. The earlier report showing the potential of decanoic acid vesicles to transition into coacervate droplets^[56]^ prompted us to question whether fatty acids having chain lengths shorter than decanoic acid be characterized for their potential to form coacervates. These also are the ones that have been poorly explored as compartment forming entities. In this context, the focus of this study was to come up with a short chain fatty acid-based coacervate system that is more stable than the two previously reported systems, which could withstand prebiotically pertinent selection pressures and that are functionally significant in the context of origins of life. Furthermore, we introduced compositional heterogeneity in our amphiphile-based coacervate system, which has been previously reported only for vesicles.^[6,13]^

To the best of our knowledge, we report for the first time a coacervate system composed of a binary mixture of short chain fatty acid and its alcohol derivative, along with the molecule tyramine (NA-NOH/Tyra). The coacervate droplets were found to be stable from pH 7-9. pH values above 9 were not considered since we intended to demonstrate a system that would support encapsulation of RNA, which gets hydrolyzed quickly in alkaline pHs. Furthermore, replacing dopamine with tyramine substantially increased the stability of the droplets. While the NA-NOH/DPA droplets could be observed only till 6 hours, the droplets formed using the NA-NOH/Tyra system could be observed for up to 8 days. It is important to note that the NA-NOH/Tyra droplets do coalesce, but readily reform upon mixing. The increased stability of NA-NOH/Tyra droplets over a long period of time does not refer to the droplets not coalescing (which does happen), but rather alluded to their ability to reform upon mixing after the reported time periods. A previous study showed that the concentration of MgCl_2_ inside the coacervates was orders of magnitude higher than the bulk solution,^[30]^ Given this, we characterized the system for this and other prebiotically relevant selection pressures. We found that NA-NOH/Tyra droplets could tolerate up to 20 mM of MgCl_2_ and 600 mM NaCl. Next, the temperature stability of coacervate droplets was evaluated and it was found that the droplets were forming until 60°C. This, coincidentally, is the proposed temperature of Darwin’s warm little pond, further underscoring the prebiotic relevance of this coacervate system.

Finally, we checked whether the system could act as a compartment to sequester biologically relevant molecules. In this context, the sequestration of GFP was found to be much better than that of the short RNA. This effect can be attributed to the potential of the protein to facilitate more interactions with the components of coacervates than RNA, through various functional groups that it contains. Moreover, as mentioned earlier, RNA being negatively charged, could experience more repulsion from the negatively charged fatty acids present in the coacervate droplets. However, we observed enhanced sequestration of RNA with the addition of positively charged amino acids, such as lysine and arginine. Our results show that the RNA sequestration could be brought about inside the coacervates by simple monomeric amino acids that would have been present as co-solutes. This implies that even relatively small co-solutes present in the prebiotic soup could have potentially altered the physicochemical properties of the coacervate droplets. This is especially appealing given that most previous coacervate research showed that such RNA sequestration necessarily needed coacervates composed of higher molecular weight cationic peptides or complex polymers.

The FRAP experiments involving the fatty acid coacervates, show that they could facilitate diffusion of molecules in the interior, suggesting the feasibility of carrying out prebiotically pertinent chemical reactions within these compartments. Towards this, when a reaction involving template-directed nonenzymatic replication of RNA was carried out in the coacervate phase, we observed extension of the primer, suggesting the compatibility of the system to prebiotically important reactions. However, it is likely that the sequestered biomolecules could also be getting exchanged with the bulk solution by diffusion as coacervates are an open system unlike membrane-bound vesicles.^[57]^ This then poses a question of whether fatty acid-based coacervates could be encapsulated by a semipermeable lipid bilayer to make them relevant in this specific context.^[58]^ Such an assembly could help with limiting the diffusion of entrapped molecules into the bulk solution while also allowing the ready diffusion within the coacervate interior. Furthermore, this work also shows how the chemical diversity of amphiphiles enhance the stability of the resultant coacervates. This invokes the importance of systematically characterizing other SCAs, including those that have been recently described as vesicle forming protoamphiphiles (e.g.N-acyl amino acids) to delineate even more robust fatty acid coacervate systems with implications for fabricating compartments with more complex functionalities.^[59,60]^

In conclusion, this work just scratches at the surface of delineating coacervate systems using prebiotically available SCA-based systems, especially on the shorter end of the chain length spectrum. Eventually, further studies are required to delve deeper into discovering the role of compositional heterogeneity in prebiotically relevant coacervate systems. This would eventually allow us to characterize SCA-based coacervate forming composite systems, allowing for effectively modulating their physicochemical properties and fabricating complex cell-like entities with multiple kinds of encapsulated compartments. For instance, compositionally heterogenous coacervates would differ in their microenvironment and this could potentially influence differential partitioning of molecules within their interior. By extension, compositionally diverse protocellular populations could potentially harbor various metabolic reactions within them, setting the stage for evolving into more complex protocells. Lastly, exploring the SCA chemical space will allow for tweaking the constituents of the coacervates that would permit the modulation of their microenvironment in different ways. Notably, this could also pave the way to develop novel delivery systems that can carry chemically distinct cargo molecules, finding relevant use both in basic and applied synthetic biology applications.

## Materials and methods

### Materials

Nonanoic acid (N5502-25G), 1-nonanol (131210-100ML), dopamine hydrochloride (PHR1090-1G) and tyramine hydrochloride (T2879-25G) were purchased from Sigma-Aldrich (Bangalore). Tris base was procured from Himedia. MgCl_2_ hexahydrate was procured from Qualigens. NaCl was purchased from Sigma-Aldrich (Bangalore). Cy3-labelled RNA (for sequestration) was obtained from Sigma (Bangalore). Glass slides and coverslips (18 mm × 18 mm) were obtained from Himedia. Adenoside-5’-phosphorimidazolides (ImpA) was purchased from GLSynthesis Inc. The RNA primer used here was terminated with a 3’-amino-2’,3’-dideoxynucleotide (Amino G primer) and was acquired from Keck laboratory, Yale, USA. It was gel purified before use. The template was purchased from Sigma-Aldrich (Bangalore) and it was used without any further purification. All other reagents (analytical grade) were purchased from Sigma-Aldrich.

Sequences of the RNAs used in the study are as follows:

1. RNA used for sequestration experiments: (Cy3) 5’ GG GAU UAA UAC GAC UCA CUG G
2. Amino G Primer: (Cy3) 5’ GG GAU UAA UAC GAC UCA CUG-NH_2_
3. Template: 5’ AGU GAU CU**U** CAG UGA GUC GUA UUA AUC CC (The nucleotide mentioned in bold is the templating base.)

## Methods

### 1. Preparation of Tyramine stock

A 500 mM Tyramine stock was prepared by dissolving tyramine hydrochloride in 200 mM Tris, pH 8. The solution was filtered through 0.22 μm syringe filter and the stock was stored in -40°C.

### 2. Preparation of NA-NOH/Tyra coacervate droplets

Nonanoic acid and nonanol were mixed in 10:1 ratio in a microcentrifuge tube at room temperature, vortexed to ensure proper mixing, and briefly centrifuged using a benchtop centrifuge to avoid sticking of the contents to the walls of the tube. Then 200 mM Tris buffer of pH 8 was added to the above mixture and vortexed vigorously to prepare 150 mM NA-NOH vesicle stock. While preparing NA-NOH/Tyra droplets, NA-NOH vesicles were added at desired concentration to the diluent (200 mM Tris buffer with pH 8) and were mixed, and the desired concentration of tyramine was then added. As soon as tyramine was added to the vesicles, the sample turned turbid, indicating the transition into coacervate droplets. It was then thoroughly mixed by gentle pipetting.

### 3. Slide preparation and microscopy

Glass slides were first rinsed with water and cleaned thoroughly, and were kept soaked in 1M HCl overnight. They were then rinsed with water extensively to remove any traces of the acid. For imaging, the coacervate sample was applied onto the clean glass slide, a coverslip was placed on it and was immediately sealed. Olympus BX63 system was used for acquiring DIC images (using 20X/0.50 and 100X/1.4 objectives) while Zeiss LSM 710 system was used for acquiring confocal images and for performing FRAP (63X/1.4 objective). All images were analyzed using FIJI (FIJI Is Just ImageJ) software.

### 4. Optical density (OD) measurements

To carry out turbidity assays, 100 μL coacervate sample was prepared and was gently mixed by pipetting. 80 μL of the sample was then dispensed in a 96 well plate for measuring the OD at 600 nm. The samples in the plate were read using Perkin Elmer EnSight plate reader.

### 5. Selection Pressure experiments

All these experiments were carried out on 60/100 NA-NOH/Tyra coacervates.

#### i) Salt tolerance

MgCl_2_ or NaCl was added to 100 μL of preformed coacervates at the desired concentration. The sample was mixed well and was imaged using DIC microscopy and checked for OD at 600 nm using plate reader. Addition of MgCl_2_ above 20 mM concentration resulted in the formation of precipitates in the sample as soon as the sample was mixed.

#### ii) Temperature tolerance

100 μL coacervate sample was prepared and was incubated at the desired temperatures for 30 minutes. They were then immediately imaged using DIC microscopy and checked for OD at 600 nm using plate reader.

### 6. Fluorescence recovery after photobleaching (FRAP)

FRAP experiments were performed by exciting the sequestered GFP within the droplets at 488 nm. The diameter of the droplets that was selected for performing FRAP ranged between 5 to 10 μm. A sequence of 10 images was taken as pre-bleach images. For bleaching, a region of interest (ROI) having an area of 1 μm^2^ was placed at the center of the droplet and it was bleached using 100% of 488 nm and 405 nm laser power, and the fluorescence intensity change in the bleached ROI was recorded by checking the emission between 493 to 797 nm. To analyze the FRAP data, ROIs of areas 1μm^2^ were used to measure the average florescence intensity of the bleached spot, the reference spot (a site away from the bleached spot on the same droplet), and that of the background (outside the droplet). Average fluorescence intensities of the aforementioned ROIs were also measured for all the pre-bleach images. As reported earlier, double normalization was carried out by using the following equation:

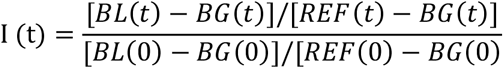

Normalized fluorescence intensities were then plotted against time, and the recovery curve was fitted assuming single exponential recovery kinetics, with the following equation:

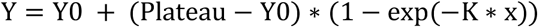

Where, Y is the fluorescence intensity at time t, Y0 is the fluorescence intensity at time 0, plateau is the fluorescence intensity at infinite times, and K is the rate constant, expressed as reciprocal of time units on X axis. Analysis was carried out in GraphPad Prism 10, and values for half-time of recovery were obtained for all the thirty droplets analyzed across the three experimental replicates. Furthermore, apparent diffusion coefficient was calculated using the following equation:^[53]^

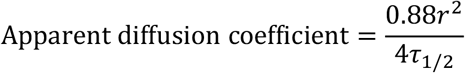

where r is the radius of the bleached area, τ_1/2_ is the half-time of recovery of fluorescence.

### 7. Laurdan Generalized Polarization (GP)

Laurdan was added at a final concentration of 5 μM to the coacervate (or vesicle) sample and the sample was incubated at room temperature for 5 mins. Laurdan emission was recorded from 400 nm to 600 nm, and GP value was calculated using the following equation:

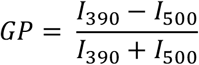

where I_390_ and I_500_ are the fluorescence intensities at 390 and 500 nm, respectively.

### 8. Confocal microscopy for RNA sequestration

Cy3 labelled RNA (5 μM) was added to 60 mM NA-NOH vesicles in 200 mM Tris, pH 8 and the sample was mixed properly. This was followed by inducing coacervation by adding tyramine at a final concentration of 100 mM. The sample was incubated for 5 mins before imaging. The sample on the glass slide was excited using 561 nm laser. We observed increased fluorescence in the droplets with the addition of lysine or arginine at the time of standardizing the experiment. Therefore, to stay below saturation in all the cases and be able to quantify images, we first imaged samples that contained arginine and standardized the microscope settings, mainly the digital gain and the laser power. After confirming that there was no saturation in any image, the rest of the samples containing no amino acid (control) were imaged under exactly identical settings. All the images were analyzed using particle analysis in FIJI. While analyzing the particles, we excluded particles below 3 μm. We recorded the integrated densities of the droplets and the mean intensity of the background. The background intensity was then subtracted from the integrated density values for the droplets and plotted these values on the Y axis. Kruskal-Wallis test for performed on the dataset by using GraphPad Prism 10.

### 9. Template-directed nonenzymatic replication of RNA in the coacervate phase

A previously standardized reaction scheme was used to perform the template-directed nonenzymatic replication reaction in the coacervate phase.^[61–63]^ Specifically, we first added the primer (0.325 μM) and template (1.3 μM) in 200 mM Tris at pH 8 and heated the sample to 90 °C for 5 mins. This was followed by flash cooling on ice and centrifuging the sample briefly. Subsequently, NA-NOH vesicles (60 mM), amino acid (25 mM Lys or Arg) and tyramine (100 mM) were added and the sample was incubated for 5 mins. The control reaction had no amino acid added to the reaction. Following this, imidazole activated nucleotide (ImpA) was added at a final concentration of 10 mM and the sample was immediately centrifuged at 14000 rcf for 2.5 mins at 15 °C. Coacervate phase was then separated from the bulk phase and was incubated at 25 °C for 2 hrs. This was followed by the addition of 7 μL TBE buffer containing 8 M urea and freezing the samples at -40 °C. Before loading the samples on 22 % denaturing urea PAGE, non-labelled RNA having an exactly identical sequence as that of the primer was added in 50 times excess when compared to the primer. The loading dye was added and this was followed by heating the sample at 90 °C for 5 mins and flash cooling on ice.^[61–63]^ For zero time point, a control reaction was set up where the coacervate phase was separated and immediately frozen as mentioned earlier albeit without incubating.

## Author Contributions

GMP conceived the project, GMP and SR designed the study. GMP carried out most of the experiments. VS helped with initial standardization of the coacervate experiments. GMP and SR analyzed the data. GMP and SR wrote the manuscript.

## Acknowledgements

SR acknowledges research support from IISER Pune, Department of Science and Technology’s Science and Engineering Research Board (SERB) (Grant No. CRG/2021/001851) and Department of Biotechnology, Govt. of India (Grant No. BT/PR43193/BRB/10/2006/2021). The authors acknowledge the microscopy facility at IISER Pune and wish to extend their special thanks to Vijay Vittal for his help with microscopy data acquisition. GMP wishes to thank the Department of Biotechnology (DBT), Government of India for fellowship support. The authors would like to thank the COoL lab members, especially Daksh Barad, for their inputs.

